# 5hmC enhances PARP trapping and restores PARP inhibitor sensitivity in chemoresistant BRCA1/2-deficient cells

**DOI:** 10.1101/2024.10.25.620335

**Authors:** Suhas S. Kharat, Arun P. Mishra, Satheesh K. Sengodan, Dillon Dierman, Stephen D. Fox, Shyam K. Sharan

## Abstract

Mutations in *BRCA1* and *BRCA2* genes are the leading cause of hereditary breast and ovarian cancer. *BRCA1/2*-mutant cells are defective in repairing damaged DNA by homologous recombination and are characterized by hypersensitivity to PARP inhibitors. PARP inhibitors can trap PARP proteins on the chromatin, a mechanism that can contribute to the death of BRCA1/2-deficient cells. The FDA has approved multiple PARP inhibitors for the treatment of metastatic breast and ovarian cancers, but despite the success of PARP inhibitors in treating *BRCA1/2*-mutant cancers, drug resistance is a major challenge. Here, we report that 5hmC enhances PARP1 trapping on the chromatin in olaparib-treated cells. Elevated PARP trapping generates replication gaps, leading to the restoration of PARP inhibitor sensitivity in chemoresistant BRCA1/2-deficient cells. Our findings suggest that combining 5hmC with olaparib can restore the sensitivity of chemoresistant BRCA1/2-deficient cells.

## Introduction

Maintenance of genomic integrity and stability is a vital function of the cellular DNA damage response (DDR), a signaling cascade activated in response to various forms of genotoxic stress. DDR consists of multiple pathways and cell cycle checkpoints that allow the cells to sense and repair the damaged DNA. A defect in the DDR results in the accumulation of unrepaired DNA damage, such as single-stranded breaks (SSB) and double-stranded breaks (DSB). SSBs are repaired by the canonical single-strand break repair pathway (SSBR) [1]. Multiple proteins are required for the repair of SSBs. Among these, the poly (ADP-ribose) polymerase (PARP) family of proteins plays an important role [2, 3]. DSBs are repaired by either the non-homologous end joining (NHEJ) pathway or homologous recombination repair (HR) [4, 5]. In the NHEJ pathway, DSBs are repaired by ligation of broken ends with less accuracy, making NHEJ error prone. In contrast, the HR pathway repairs DSBs with high fidelity. Among the proteins with an important role in HR-mediated DSB repair are BRCA1, BRCA2, and PALB2. These are all well-established tumor suppressors associated with an increased risk of breast, ovarian, and other cancers [6, 7].

The research community has exploited the essential role of DNA repair proteins for cell viability to develop novel chemotherapeutics. PARP inhibitors (PARPi), one class of such chemotherapeutics, blocks the repair of SSBs, resulting in replication fork stalling and replication-associated DSB and subsequently making the cells dependent on HR-mediated DNA repair for survival [3]. Cells deficient in BRCA1 or BRCA2, proteins essential for HR, are known to be sensitive to PARPi. This forms the basis of PARPi-induced synthetic lethality in such cells. Through the synthetic lethality mechanism, a PARPi blocks the repair of DNA SSBs in tumor cells with an HR defect, causing them to accumulate a large number of DNA DSBs. This induces killing of BRCA1/2-deficient tumor cells and, thus, generates significant anti-tumor activity in patients [8, 9]. In a Phase 1 clinical trial of the PARPi olaparib conducted in *BRCA1* and *BRCA2* germline mutation carriers with advanced solid tumors, 63% of treated patients demonstrated clinical benefit [10]. Many clinical trials have extended their studies, with promising results in other tumors exhibiting a characteristic phenotype of HR deficiency known as BRCAness: triple-negative breast cancers, advanced prostate cancers, and ovarian cancers [11, 12]. Olaparib, rucaparib, and niraparib are currently Food and Drug Administration (FDA)-approved PARP inhibitors for patients with platinum sensitivity and recurrent ovarian cancer [11, 13, 14]. Olaparib and talazoparib also have received FDA approval for metastatic breast cancer with a germline *BRCA* mutation [15, 16]. HR is regulated by many genes besides *BRCA1/2*, as tumors proficient in BRCA1/2 are also reported to exhibit sensitivity to PARPi [17, 18]. Thus, in principle, mutations in different genes involved in the HR pathway could bear similar consequences to mutations in *BRCA1/2*. The use of PARPi has accordingly expanded to other genes that result in BRCAness phenotype. For instance, PARPi have improved survival of prostate cancer patients with mutations in *PALB2, RAD51C/D,* and *CHEK2* [19].

Despite the success of PARPi in the clinic, tumors in more than 40% of patients have exhibited resistance to PARPi chemotherapy, which poses a major challenge [20, 21]. Several mechanisms are responsible for PARPi resistance. Secondary mutations in *BRCA1/2* that restore HR are well-known contributing factors [22–24]. Loss of several accessory proteins that are associated with DNA end resection, such as 53BP1, RIF1, and members of the shieldin complex (including REV7), has been demonstrated to restore HR in BRCA1-deficient cells [25–28]. PARPi resistance has been observed in HR-deficient BRCA2-mutant (p.L2510P) mouse embryonic fibroblasts (MEFs) due to homologous DNA present in the vicinity of replication-induced DSBs, which led to the formation of RAD51 foci [29]. PARPi resistance is also observed when stalled replication forks (RF) can be protected in BRCA1/2-deficient cells. BRCA1/2 are required for maintaining the stability of stalled RFs [30, 31] from degradation by nucleases such as MRE11 and MUS81 [32]. Loss of EZH2 and PTIP in BRCA2-deficient cells abolishes recruitment of MRE11 at stalled RF and results in fork stabilization [33, 34]. Loss of TET2 protects RF from APE1-endonuclease-mediated degradation in BRCA2-deficient cells and confers PARPi resistance [35]. Increased drug efflux due to overexpression of ABC transporters, such as P-glycoproteins (PgP), has been demonstrated to confer PARPi resistance [36]. PARPi lethality in BRCA1/2-deficient cells varies and largely depends on the ability of the inhibitors to bind and trap PARP proteins on the chromatin. Talazoparib and olaparib are excellent PARP trappers and are therefore more effective in killing BRCA1/2-deficient cells than veliparib [37]. However, the presence of PARP also influences sensitivity to PARPi, and loss of or decrease in PARP levels can confer PARPi resistance in human cancer cells [38, 39].

Since resistance to PARPi is frequently observed in patients’ cancers, novel approaches are being developed to overcome it. Novobiocin (NVB), an antibiotic that inhibits the activity of POLQ, has been shown to restore sensitivity in 53BP1-BRCA1 double-knockout cells [40]. Cell cycle signaling inhibitors have been investigated to overcome PARPi resistance in BRCA1-deficient tumors. Some ovarian cancer cells that are defective in HR and RF protection acquire resistance to PARPi or cisplatin due to elevated ATR-CHK1 activity. Combining ATR inhibitors restored PARPi sensitivity in these cells [41]. It has also been demonstrated that loss of 53BP1-RIF-REV7-shieldin complex impairs the NHEJ pathway and restores HR in BRCA1-deficient cells. Loss of shieldin complex renders tumor cells sensitive to radiation [42]. It remains to be seen whether this vulnerability can be utilized to treat such BRCA1-deficient tumors with acquired resistance to PARPi.

Vitamin C (VitC) has been evaluated as a possible anti-cancer agent [43, 44]. Supportive evidence from a plethora of preclinical and clinical studies suggests that VitC is effective either as a monotherapy or in conjunction with conventional chemotherapies. A high dose of VitC has been shown to suppress growth of breast and ovarian cancer cells [45, 46]. A combination of high doses of VitC and PARPi synergistically inhibits the growth of BRCA1/2-proficient cells by enhancing DNA DSBs. Based on these observations, use of VitC has been recommended to enhance the efficacy of PARPi [47], and findings from a clinical trial suggest that a high dose of VitC could be a plausible additional therapy with PARPi in HR deficiency [48]. Moreover, a combinatorial therapy of high-dose VitC and PARPi has also been reported for treatment of Ewing sarcoma [49] and acute myeloid leukemia (AML) [50].

VitC is a co-factor for a range of enzymes, including Fe(II)- and α-ketoglutarate dioxygenases (α-KGDD), such as TET dioxygenases. Furthermore, VitC enhances the activity of TET proteins and promotes demethylation leading to genomic instability [51]. TET proteins (TET1/2/3) are members of a family of α-KGDDs that catalyze iterative oxidation of 5mC to 5-hydroxymethylcytosine (5hmC), 5-formylcytosine (5fC), and 5-carboxycytosine (5caC) [52]. Due to the low affinity of 5hmC to TET proteins, it is the most stable modification and constitutes 10% of total 5mC [53]. 5hmC is reported to be greatly reduced or suppressed in multiple cancers [54]. Its role remains elusive, but certain findings suggest that it acts as an epigenetic marker for DNA damage and may recruit specific readers for direct dynamic remodeling and organization of chromatin structures. In addition, increased accumulation of 5hmC can elicit chromosome segregation and RF degradation [35, 55].

In the present study, we have described the role of 5hmC and VitC in restoring PARPi sensitivity in BRCA1/2-deficient chemoresistant cells. We used HR-restored BRCA1-53BP1 double-knockout cells and RF-protected CHD4/PTIP-BRCA2 double-knockdown cells as chemoresistant models. Our findings revealed that combining 5hmC and VitC with olaparib led to increased PARP trapping in chemoresistant BRCA1/2-deficient cells. 5hmC/VitC-dependent PARP trapping generated replication gaps and restored olaparib sensitivity.

## RESULTS

### VitC and 5hmC enhance efficacy of PARP inhibitor in BRCA1/2-deficient cells

We have previously reported that enhancing TET activity by its cofactor VitC results in the degradation of stalled RFs by APE1 endonuclease [35]. Furthermore, VitC enhances the degradation of stalled RFs in BRCA2-deficient cells. Therefore, we hypothesized that VitC can enhance the sensitivity of these cells when combined with PARPi and potentially restore PARPi sensitivity in chemoresistant cells. We tested the effect of VitC combined with olaparib in an isogenic pair of human UWB1.289-BRCA1WT/UWB1.289-BRCA1KO and DLD1-BRCA2WT/DLD-BRCA2KO (hereafter referred to as UWB1.289-WT/BRCA1KO and DLD1-WT/BRCA2KO) cells. Clonogenic and XTT cell proliferation assays revealed that the combined treatment did not significantly affect viability of UWB1.289-WT and DLD1-WT cells. However, viability of UWB1.289-BRCA1KO (Figure S1A-C) and DLD1-BRCA2KO (Figure S1D-F) cells was significantly reduced when treated with VitC combined with olaparib compared to olaparib alone. A similar increase in VitC-dependent olaparib sensitivity was also observed in MEF-BRCA1KO (Figure S1G-I). To compare the effects of VitC and olaparib between BRCA1- and BRCA2-deficient cells, we performed siRNA-mediated knockdown of *BRCA1* and *BRCA2* in DLD1-WT cells (Figure S1J). The combined VitC-olaparib treatment decreased viability of siBRCA1 and siBRCA2 cells but has no effect on siControl cells (Figure S1K-L). VitC treatment alone had significantly reduced the viability of all tested BRCA1/2-deficient cells and did not show a synergistic effect when combined with PARPi. The PARPi-independent effects of VitC on *BRCA1/*2-KO cells are corroborated by previous reports. For instance, multiple types of cancer cells, including breast cancer, have been reported to be sensitive to VitC because of its ability to increase global 5hmC [56, 57]. These observations suggests that effect of VitC on PARPi is additive rather than synergistic.

VitC at lower concentrations is known to act as an antioxidant exerting protection against reactive oxygen species (ROS) levels in normal cells. However, at higher concentrations, VitC acts as a pro-oxidant and helps in preferentially accumulating cytotoxic ROS levels to exert anti-tumorigenic effects on cancer cells. The presence of metal ions, such as iron, affects the switch between these antioxidant and pro-oxidant roles [58, 59]. To determine mechanism of VitC sensitivity in BRCA1/2 KO cells, we examined ROS levels in UWB1.289-WT/BRCA1KO and DLD1-WT/BRCA2KO cells after treatment with VitC and olaparib individually and together. ROS levels in both pairs of cell lines were found to be unaltered after VitC and/or olaparib treatments (Figure S2A-B).

Next, we evaluated the effect of VitC on the activity of TET proteins by quantitating global 5mC and 5hmC levels by mass spectrometry. Our results revealed a global increase in 5hmC levels in response to VitC treatment in UWB1.289-WT/BRCA1KO (Figure S2C-D) and DLD1-WT/BRCA2KO cells (Figure S2E-F). TET proteins require PARylation for their activity, and PARP inhibition reduces PARylation of TET1, causing reduced activity and a global decrease in 5hmC levels [60]. Consistent with this, we found PARP inhibition by olaparib reduced global 5hmC levels in UWB1.289-WT/BRCA1KO (Figure S2G) and DLD1-WT/BRCA2KO cells (Figure S2H). However, combined treatment with VitC and olaparib compensated for the loss of global 5hmC levels and even significantly increased 5hmC levels in UWB1.289-BRCA1KO (Figure S2C) and DLD1-BRCA2KO (Figure S2E) cells compared to their respective isogenic WT controls. Since VitC enhances TET activity and sensitivity to PARPi, we analyzed the effect of individual cytosine analogues (i.e., 5mC, 5hmC, 5fC, and 5caC), catalyzed by TET proteins, in UWB1.289-WT/BRCA1KO and DLD1-WT/BRCA2KO cells. Clonogenic and XTT drug assays revealed that 5hmC specifically enhanced sensitivity to olaparib in UWB1.289-BRCA1KO (Figure 1A-C) and DLD1-BRCA2KO cells (Figure 1D-F) when compared to their respective isogenic WT cells. Effect of 5hmC on olaparib sensitivity was synergistic because we observed a significant reduction in viability of UWB1.289-BRCA1KO (Figure 1G) and DLD1-BRCA2KO (Figure 1H) cells when increasing gradient of olaparib concentration was combined with 5hmC.

**Figure 1.**
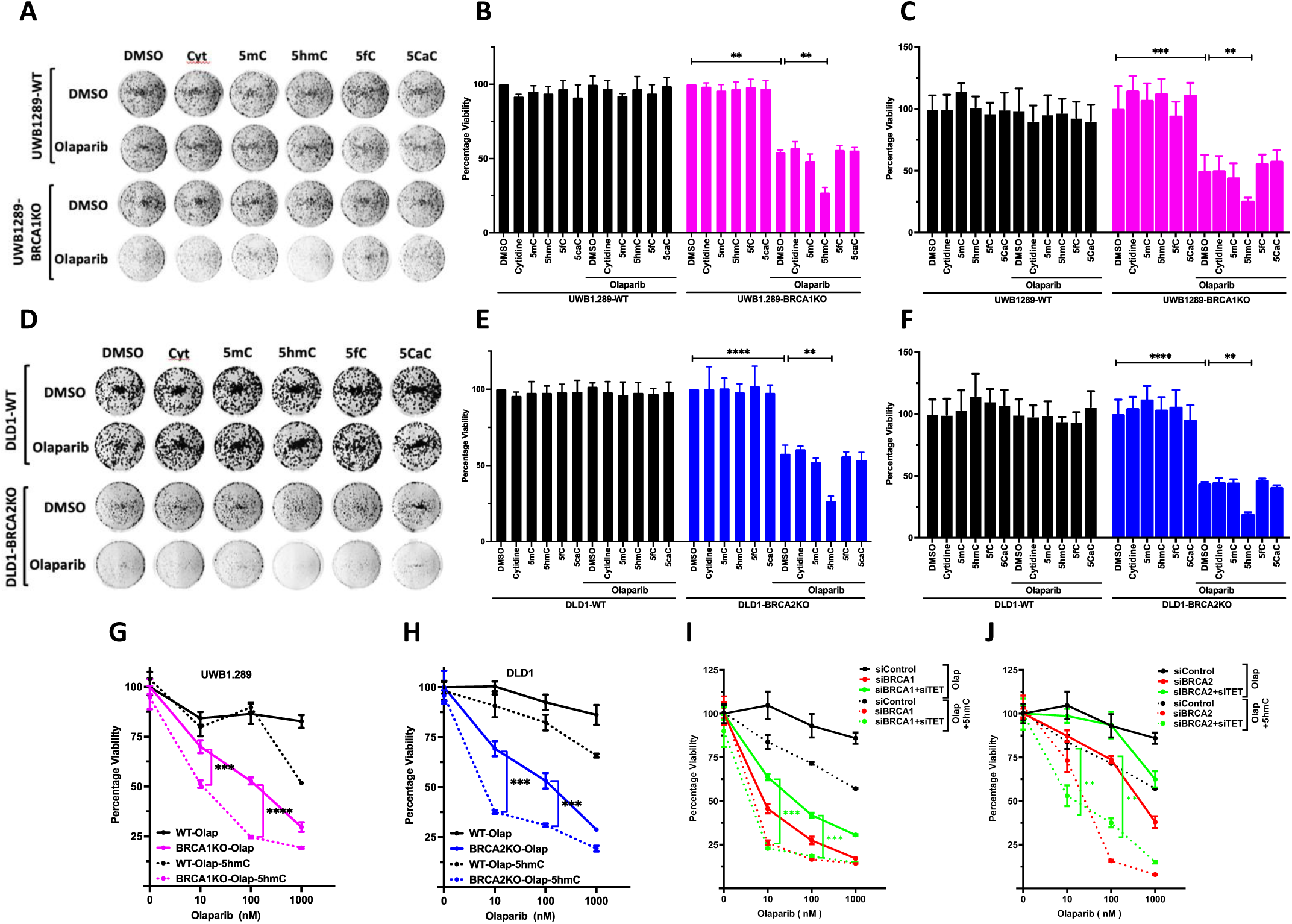
5hmC enhances PARP inhibitor sensitivity in BRCA1/2 deficient cells. **A, B, C.** Clonogenic survival assay (A), quantitation of clonogenic assay (B) and XTT survival assay (C) was performed in UWB1.289-WT/BRCA1KO cells treated with different cytosine analogues (Cytidine, 5mC, 5hmC, 5fC and 5caC) combined with olaparib (100 nM). N = 3. **D, E, F.** Clonogenic survival assay (D), quantitation of clonogenic assay (E) and XTT survival assay (F) was performed in DLD1-WT/BRCA2KO cells treated with cytosine analogues combined with olaparib (100 nM). **G, H.** XTT survival assay was performed in UWB1.289-WT/BRCA1KO (G) DLD1-WT/BRCA2KO (H) cells treated with increasing concentration of olaparib (10, 100 or 1000 nM) combined with or without 5hmC (1 μM). N = 3. **I, J.** XTT cell survival assay was performed in DLD1 cells treated with indicated drug treatments upon transient knockdown of either *BRCA1* (I) or *BRCA2* (J) combined with TET1,2,3. N = 3. Data in XTT assay is mean ± SD from 8 technical replicates. Values in XTT assay have been normalized with respective DMSO control of cell line. N = 3. Statistical analysis was performed using paired T test.

To confirm whether the effect of VitC on olaparib sensitivity was TET/5hmC dependent, we knocked down *TET1, TET2,* and *TET3* (combined knockdown refered as *siTET*) in combination with *BRCA1* or *BRCA2* in DLD1 cells (Figure S2I-J). These cells were treated with olaparib together with VitC or 5hmC; and performed XTT-based cell proliferation assay. VitC-dependent enhancement in olaparib sensitivity was abolished in *siTET-siBRCA1* and *siTET-siBRCA2* cells in comparison to *siBRCA1* or *siBRCA2* cells, respectively (Figure S2K-L). However, 5hmC was able to enhance olaparib sensitivity even in s*iTET-siBRCA1* and *siTET-siBRCA2* cells, which was comparable to the sensitivity observed in *siBRCA1* and *siBRCA2* cells, respectively (Figure 1I-J). These findings suggest that the effects of VitC are mediated by a TET-dependent increase in 5hmC levels, which can be bypassed by directly providing 5hmC to the cells.

### 5hmC restores PARPi sensitivity in chemoresistant BRCA1/2-deficient cells

We next examined whether combining 5hmC or VitC with olaparib can restore PARPi sensitivity in chemoresistant BRCA1/2-deficient cells. Loss of 53BP1 in BRCA1-deficient cells results in restoration of homologous recombination leading to PARPi resistance [26]. Therefore, we examined the effect of 5hmC or VitC on olaparib sensitivity in chemoresistant *Brca1*Δ*11;53Bp1KO* MEFs. We performed clonogenic assay as well as XTT assay to determine the effect of VitC-olaparib combined treatment. As expected, *Brca1*Δ*11* MEFs were sensitive to olaparib treatment and *Brca1*Δ*11;53Bp1KO* MEFs were resistant. However, the VitC-Olaparib combined treatment markedly affected the survival of *Brca1*Δ*11;53Bp1KO* MEFs (Figure S3A-C). VitC dependent restoration of olaparib sensitivity was further confirmed in DLD1 cells with siRNA-mediated knockdown of *BRCA1* and *53BP1* (Figure S3D-G). The loss of CHD4 and PTIP in BRCA2-deficient cells has been shown to protect stalled RF and confer resistance to PARP inhibitors. [33, 61]. To generate chemoresistant cells with RF protection, we used siRNA to knock down *CHD4* and *PTIP* in DLD1-BRCA2KO cells (Figure S3H). *siCHD4* and *siPTIP* DLD1-BRCA2KO cells exhibited resistance to olaparib when compared to siControl cells. However, viability of these cells was significantly reduced when olaparib treatment was combined with VitC (Figure S3H-K). Once again, VitC treatment alone had significantly reduced the viability of all tested chemoresistant BRCA1/2-deficient cells and did not exhibit synergistic effect when combined with PARPi. Based on the effect of VitC on TET activity and sensitivity to PARPi, we analyzed the effect of individual cytosine analogues (i.e., 5mC, 5hmC, 5fC, and 5caC), catalyzed by TET proteins, in MEF-WT, *Brca1*Δ*11*, and *Brca1*Δ*11;53Bp1KO* cells. XTT based drug assays revealed that 5hmC specifically enhanced sensitivity to olaparib in *Brca1*Δ*11* and *Brca1*Δ*11;53Bp1KO* MEFs (Figure S3L, 2A-B). Mass-spectrometry-based measurements revealed a significant global increase in 5hmC levels in WT, *Brca1*Δ*11,* and *Brca1*Δ*11;53Bp1KO* MEFs after treatment with VitC alone and VitC and olaparib together (Figure S3M-N). As observed previously, olaparib treatment alone reduced global 5hmC levels in these MEFs (Figure S3O). 5hmC-dependent restoration of olaparib sensitivity was further confirmed in DLD1 cells with siRNA-mediated knockdown of *BRCA1* and *53BP1* (Figure 2C-E). Next, we tested the effect of combined treatment of 5hmC and olaparib in chemoresistant *BRCA2-*deficient cells. Similar to VitC, olaparib sensitivity was restored in *siCHD4* and *siPTIP* DLD1-BRCA2KO cells in the presence of 5hmC (Figure 2F-H). 5hmC (1 μM) was found to be more effective in re-sensitizing PARPi-resistant cells than VitC (1 mM).

**Figure 2.**
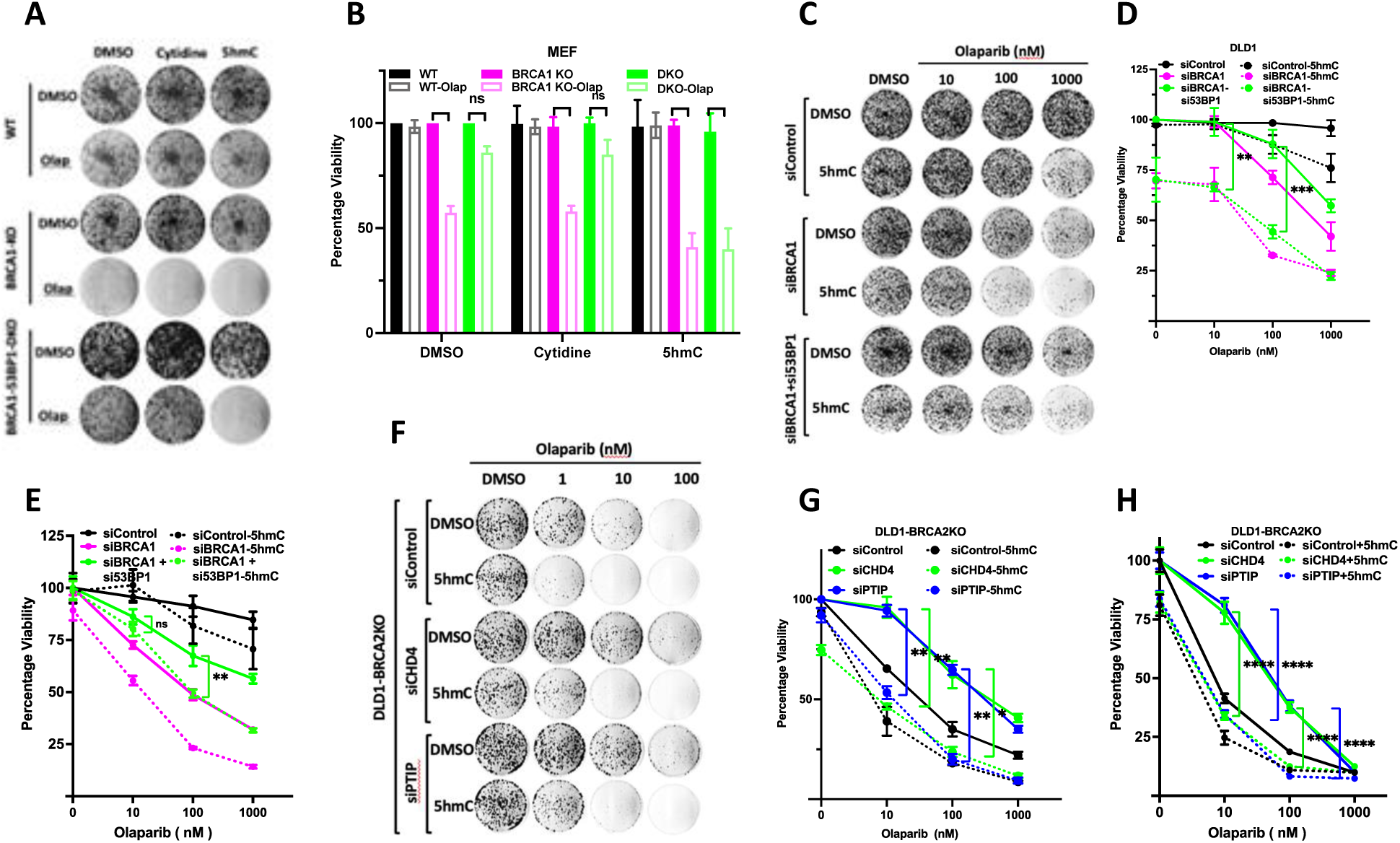
5hmC restores PARP inhibitor sensitivity in chemoresistant BRCA1/2 deficient cells. **A, B.** Clonogenic survival assay (A) and quantitation of clonogenic survival (B) was peformed in WT/*Brca1*Δ*11*/*Brca1*Δ*11;53Bp1KO* MEF cells treated with cytidine and 5hmC combined with olaparib (100 nM). N = 3. **C, D, E.** Clonogenic survival assay (C), quantitation of clonogenic assay (D) and XTT survival assay (E) was performed in DLD1-WT cells treated with indicated drug treatments upon transient knockdown of BRCA1 and 53BP1. **F, G, H.** Clonogenic survival assay (F), quantitation of clonogenic assay (G) and XTT survival assay (H) was performed in DLD1-BRCA2KO cells treated with indicated drug treatments upon transient knockdown of CHD4 and PTIP. Data in XTT assay is mean ± SD from 8 technical replicates. Values in XTT assay have been normalized with respective DMSO control of cell line. N = 2. Statistical analysis was performed using paired T test.

### 5hmC and VitC exacerbate PARP trapping in chemoresistant BRCA1/2-deficient cells

HR deficiency renders BRCA1/2 deficient cells sensitive to PARPi [62]. One of the mechanisms underlying the killing of HR-deficient cells by PARPi is the trapping of PARP proteins on the chromatin, which is toxic to the cells. We used 5hmC in combination with olaparib to examine the effect on PARP trapping in UWB1.289-BRCA1KO and DLD1-BRCA2KO cells. Cells were treated with various concentrations of 5hmC for 48 hours, followed by treatment with or without olaparib (10 μM) for 2 hours. We found 5hmC alone did not trap PARP1 in UWB1.289-BRCA1KO and DLD1-BRCA2KO cells at any concerntration that we tested. However, increased PARP1 trapping was observed in olaparib-treated UWB1.289-BRCA1KO (Figure 3A) and DLD1-BRCA2KO cells (Figure 3B) in a 5hmC-concentration-dependent manner. Similar levels of PARP1 trapping were observed in UWB1.289-BRCA1KO (Figure S4A) and DLD1-BRCA2KO cells (Figure S4B) in a VitC-concentration-dependent manner. PARP trapping is weak in olaparib treated BRCA-deficient cells in absence of genotoxic stress but is markedly increased when olaparib treatment is combined with other genotoxic agents, such as methyl methane sulfonate (MMS) [37]. We examined PARP1 trapping without MMS treatment, which would mask the effect of 5hmC/VitC-induced trapping by olaparib.

**Figure 3.**
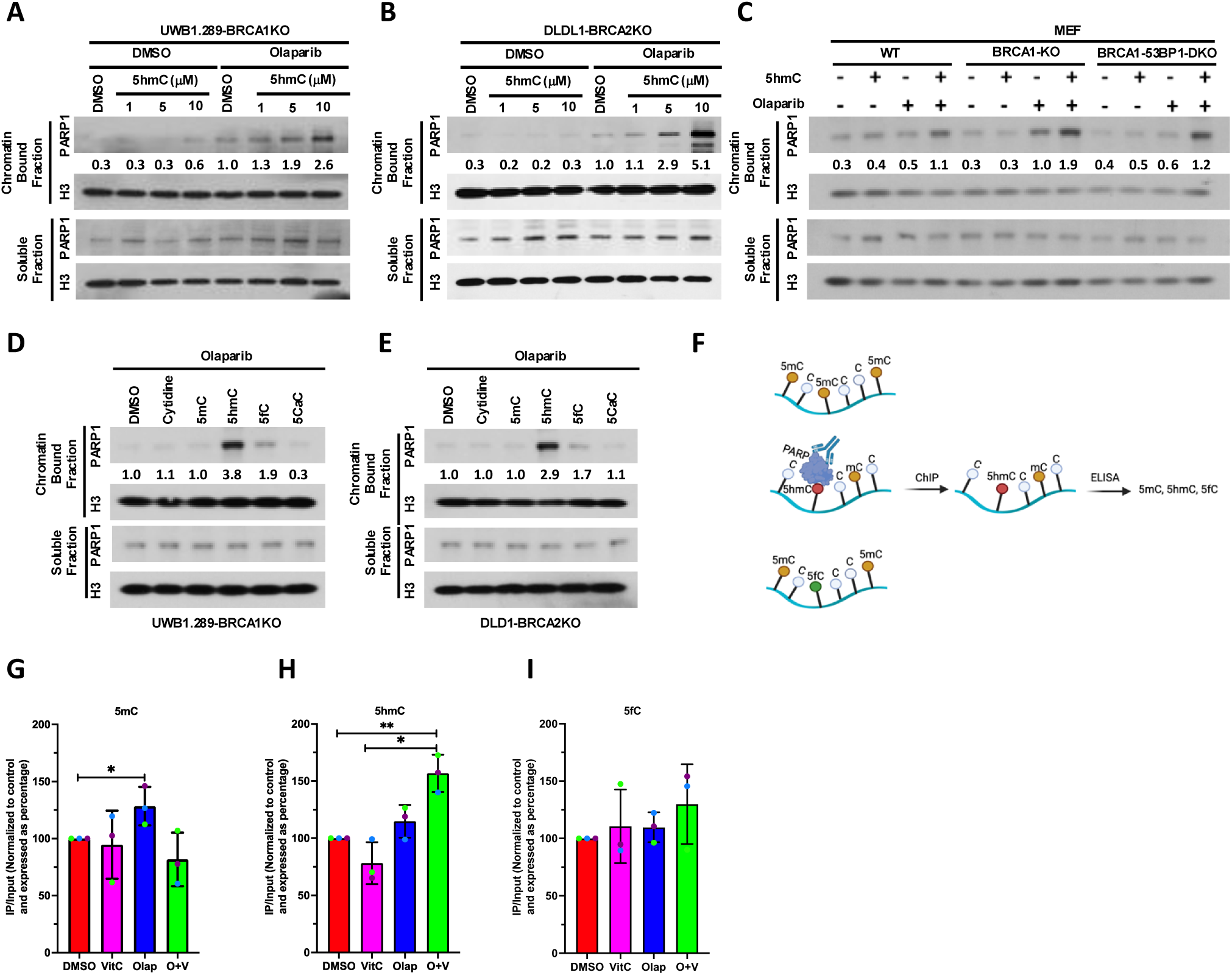
5hmC traps PARP1 in presence of olaparib in BRCA1/2 deficient cells. **A, B. C** Western blotting of nuclear soluble and chromatin-bound fractions prepared from UWB1.289-BRCA1KO (A), DLD1-BRCA2KO (B) and WT/*Brca1*Δ*11*/*Brca1*Δ*11;53Bp1KO* MEFS cells (C) upon treatment with gradient concentration of 5hmC (1, 5, 10 μM, 48 hrs) combined with or without olaparib (10 μM, 2 hrs). Blots were probed with indicated antibodies. **D, E.** Western blotting of nuclear soluble and chromatin-bound fractions prepared from UWB1.289-BRCA1KO (D) and DLD1-BRCA2KO (E) cells upon treatment with indicated cytosine analogue (10 μM, 48 hrs) combined with olaparib (10 μM, 2 hrs). Blots were probed with indicated antibodies. Band intensities for chromatin bound fractions were measured using FIJI. Chromatin bound PARP1 intensity was determined by first normalizing the values for each chromatin bound PARP1 lane to the corresponding chromatin bound H3 signal and then to same value calculated for respective cell line treated with olaparib. **F.** Schematic of ChIP-ELISA for PARP1 and 5mC, 5hmC or 5fC estimation. **G, H, I.** Detection of cytosine analogues-5mC (G), 5hmC (H) and 5fC (I) in the genomic DNA pulled down by anti PARP antibody and detected by analogue specific ELISA kit. Bars represent percent increase in 5mC level when cells were treated with DMSO, LAA, olaparib and combination of olaparib and Vitamin. All values are normalized to their repective input control.

Next, we tested the effects of 5hmC on PARP trapping in *Brca1*Δ*11* (sensitive) and *Brca1*Δ*11;53Bp1KO* (resistant) MEFs. Significantly higher levels of PARP trapping were observed in olaparib-treated *Brca1*Δ*11* MEFs compared to *Brca1*Δ*11;53Bp1KO* MEFs. However, 5hmC-olaparib combined treatment likewise significantly enhanced PARP1 trapping in *Brca1*Δ*11;53Bp1KO* MEFs, which was comparable to that observed in *Brca1*Δ*11* MEFs (Figure 3C). Similar to 5hmC, VitC-olaparib combined treatment also resulted in PARP1 trapping in *Brca1*Δ*11;53Bp1KO* MEFs comparable to *Brca1*Δ*11* MEFs (Figure S4C). We also compared PARP1 trapping induced by cytosine analogues (5mC, 5hmC, 5fC, and 5caC) in UWB1.289-BRCA1KO and DLD1-BRCA2KO cells. Efficient PARP1 trapping was observed in both cell lines after combined treatment of 5hmC and olaparib (Figure 3D-E).

To corroborate these findings, we performed ChIP-ELISA (Figure 3F) in DLD1-BRCA2KO cells to measure the levels of 5mC, 5hmC and 5fC in PARP trapped genomic DNA (Figure 3G-I). We observed significant PARP trapping in genomic DNA fragments enriched in 5hmC in cells with combined treatment of VitC and olaparib (Figure 3H). However, significant PARP trapping was not observed in genomic DNA fragments enriched in 5mC and 5fC under combined treatment of VitC and olaparib (Figure 3G, 3I). We found olaparib treatment resulted in mild but significant enrichment of 5mC. It does not necessarily suggest that PARP gets trapped at 5mC sites. Based on the input values, 5mC is the predominant analog compared to 5hmC and 5fC. Therefore, we speculate that PARP trapping occurs at 5hmC sites, but the pulled-down sonicated DNA fragments (150-400bp) possess more 5mC sites in the vicinity of 5hmC sites, when the cells are treated with olaparib.

### Exposure to VitC and 5hmC induces replication gaps and DNA damage

Genotoxic drugs induce replication stress, usually characterized by global replication slowdown or stalling. Replication restarts after the repair of damaged DNA. In case of replication re-initiation blockage, cancer cells employ DNA damage tolerance pathways, such as firing new origins or repriming of replication initiation downstream from the stress site, resulting in under-replicated DNA and replication gaps [63]. The generation of replication gaps can cause replication catastrophe and hence can be leveraged to enhance the efficacy of genotoxic chemotherapy [64]. PARPi-sensitive cells exhibit an increase in single-stranded gaps [65–67]. To gain insight into the effect of 5hmC and VitC on the generation of replication gaps, we performed DNA fiber assays to monitor RF dynamics. 5hmC incorporation has been shown to increase PAR levels in BRCA2-deficient cells. It has been reported that 5hmC repair by PARP1 during base excision repair (BER) interfered with RF progression and contributed to genomic instability [68]. We used UWB1.289-WT/BRCA1KO and DLD1-WT/BRCA2KO cells to measure the replication track lengths after 5hmC and olaparib treatment. Replication tracks were significantly longer in olaparib-treated UWB1.289-BRCA1KO and DLD1-BRCA2KO cells relative to their isogenic WT controls. Interestingly, the combined treatment of 5hmC-olaparib resulted in relatively long replication track lengths in UWB1.289-BRCA1KO (Figure 4A-B) and DLD1-BRCA2KO cells (Figure 4C-D) compared to when the cells were treated with olaparib alone. To determine the presence of replication gaps in these long replication tracks, cells were labeled with IdU, a thymidine analogue, and then treated with S1 nuclease to digest any ssDNA in the IdU-labeled DNA. The S1 nuclease treatment reduced the track lengths in UWB1.289-BRCA1KO and DLD1-BRCA2KO cells treated with olaparib alone and with 5hmC (Figure 4A-D). These findings were also confirmed in *Brca1*Δ*11* and *Brca1*Δ*11;53Bp1KO* MEFs. We found the replication track lengths to be significantly shorter in olaparib-treated *Brca1*Δ*11;53Bp1KO* MEFs compared to the *Brca1*Δ*11* MEFs. However, in the presence of 5hmC, there was a significant increase in replication track lengths in the *Brca1*Δ*11;53Bp1KO* MEFs (Figure 4E-F). Effects of VitC were similar to 5hmC in UWB1.289-WT/BRCA1KO (Figure S5A-B), DLD1-WT/BRCA2KO (Figure S5C-D), *Brca1*Δ*11* and *Brca1*Δ*11;53Bp1KO* MEFs (Figure S5E-F).

**Figure 4.**
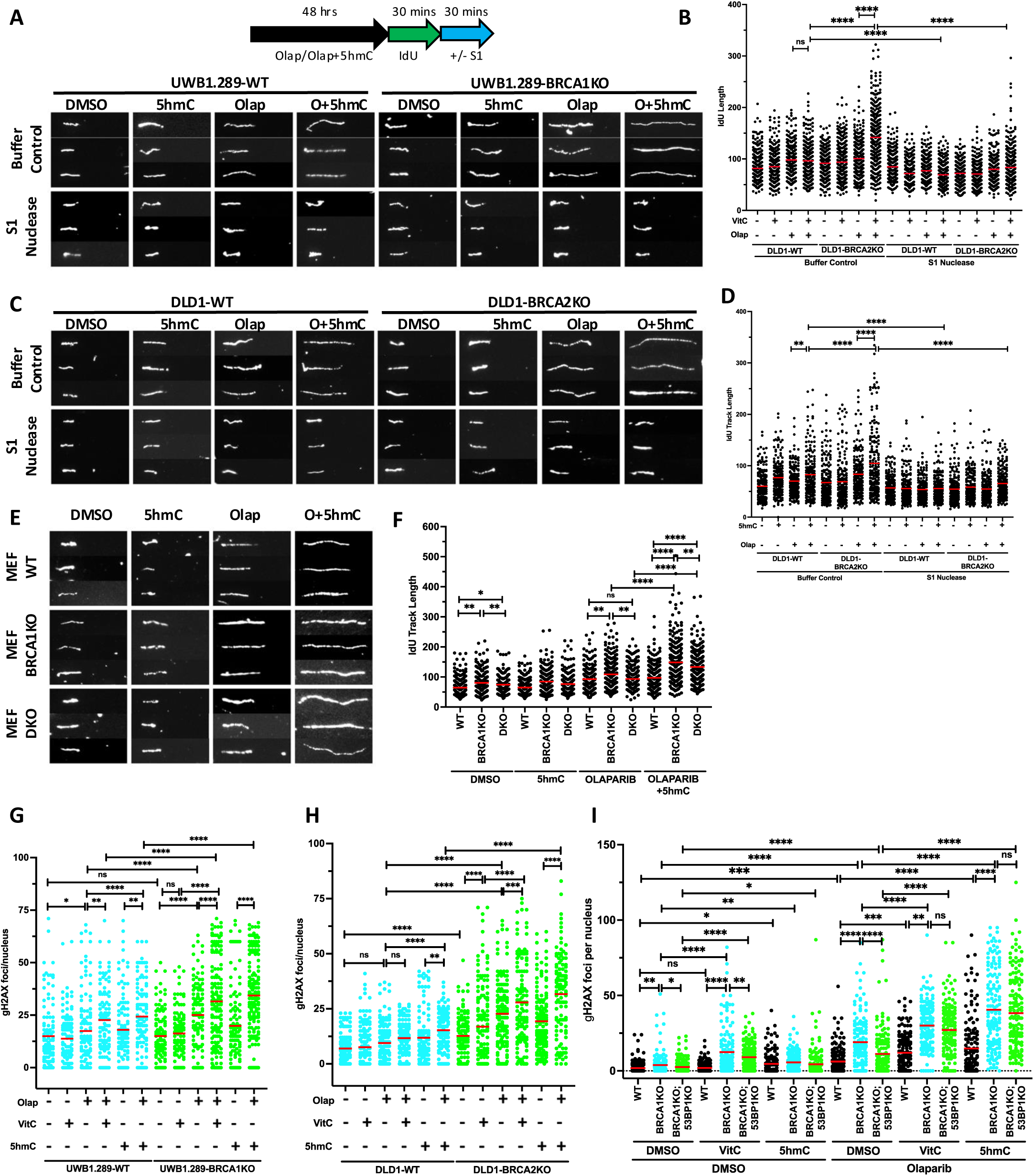
5hmC increases replication gaps and DNA damage in olaparib treated BRCA1/2 deficient cells. **A, B.** Representative DNA fibers (A), quantitation of DNA fiber length (B) for IdU tracts with or without S1 nuclease incubation in UWB1.289-WT/BRCA1KO cells upon treatment with 5hmC (1 μM, 48 hrs) and olaparib (100 nM, 48 hrs) or both. **C, D.** Representative DNA fibers (C), quantitation of DNA fiber length (D) for IdU tracts with or without S1 nuclease incubation in DLD1-WT/BRCA2KO cells upon treatment with 5hmC (1 μM, 48 hrs) and olaparib (100 nM, 48 hrs) or both. **E, F.** Representative DNA fibers (E), quantitation of DNA fiber length (F) for IdU tracts with or without S1 nuclease incubation in WT/*Brca1*Δ*11*/*Brca1*Δ*11;53Bp1KO* MEF cells upon treatment with 5hmC (1 μM, 48 hrs) and olaparib (100 nM, 48 hrs) or both. N = > 200 fibers (N = 2). Red bars represent the mean. **G, H, I.** γH2AX foci/cell quantitation after indicated drug treatments in UBW1.289-WT/BRCA1KO (G), DLD1-WT/BRCA2KO (H) and MEF-WT/*Brca1*Δ*11*/*Brca1*Δ*11;53Bp1KO* (I) cells. Nuclei count >150. N = 2. ns P ≥ 0.05, *P ≤ 0.05, **P ≤ 0.01, ***P ≤ 0.001, and ****P ≤ 0.0001 by Mann-Whitney test.

**Figure 5.**
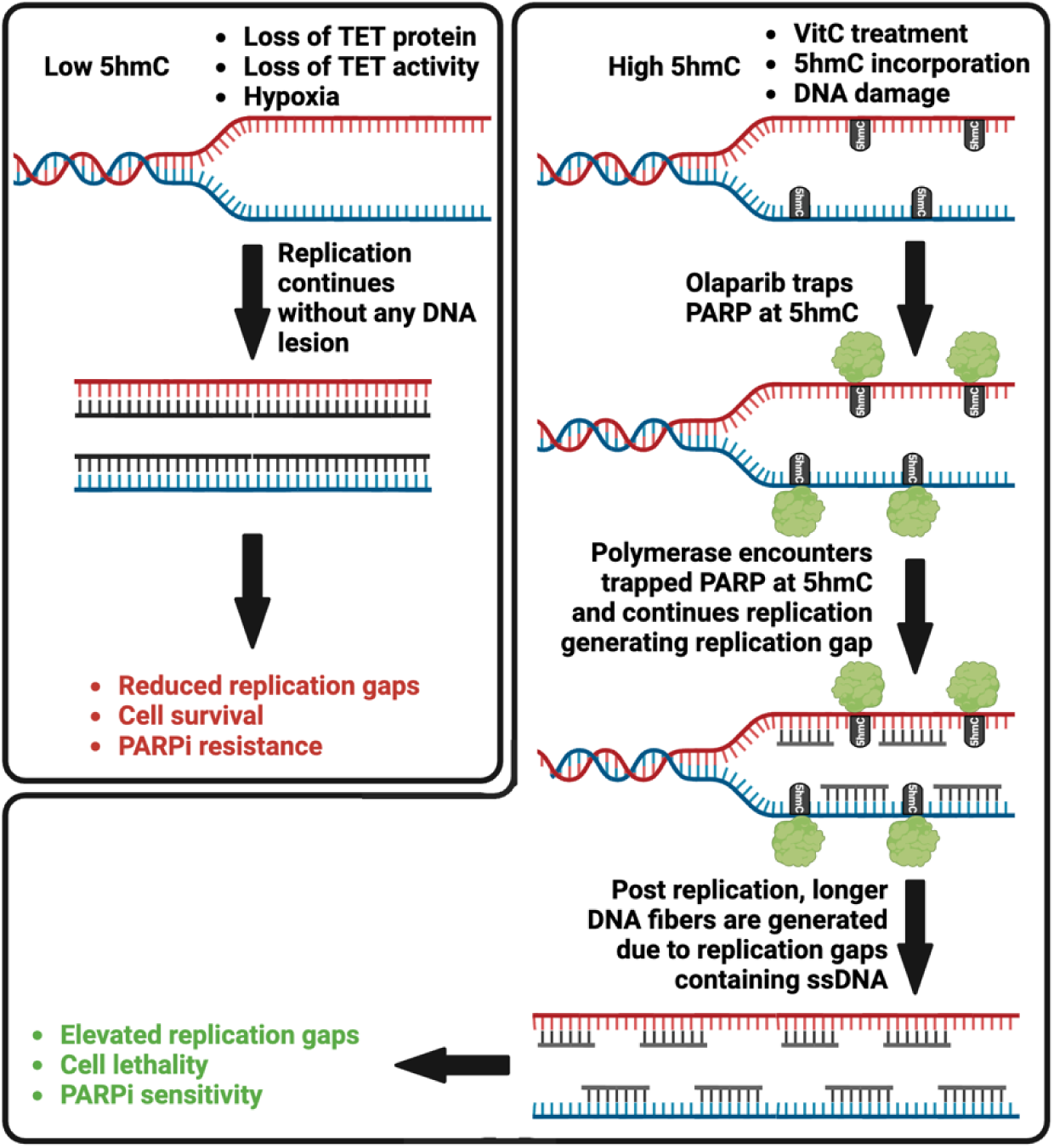
Schematic representation of VitC/5hmC dependent generation of replication gaps. Elevated 5hmC levels due to DNA damage or VitC treatment, traps PARP proteins on chromatin resulting in the generation of replication gaps. Increased replication gaps are hallmark of PARP inhibitor sensitivity and cellular lethality of BRCA1/2-deficient cells.

We examined the extent of DNA damage by quantitating γH2AX foci, a marker for DNA DSBs, in UWB1.289-WT/BRCA1KO cells, DLD1-WT/BRCA2KO cells, and WT/*Brca1*Δ*11*/*Brca1*Δ*11; 53Bp1KO* MEF cells. We found combining 5hmC or VitC with olaparib significantly increased γH2AX foci in UWB1.289-BRCA1KO (Figure 4G, S5G) and DLD1-BRCA2KO (Figure 4H, S5H) cells compared both to their respective WT controls and to treatment with olaparib alone. *Brca1*Δ*11;53Bp1KO* MEFs had significantly less γH2AX foci than *Brca1*Δ*11* MEFs after olaparib treatment, but the numbers significantly increased and were comparable to *Brca1*Δ*11* MEFs, after olaparib was combined with 5hmC or VitC (Figure 4I, S5I). Taken together, these results suggest that the combined treatment of olaparib and 5hmC or VitC induces DNA damage and replication gaps by increasing PARP1 trapping, thereby resensitizing the chemoresistant cells toward olaparib.

## Discussion

The TET proteins are known to be involved in diverse biological processes, such as epigenetic gene regulation, embryonic development, and cancer [69, 70]. The significance of the TET proteins lies in their ability to catalyze oxidation of 5mC to 5hmC, 5fC, and 5caC, which is critical for DNA demethylation. TET-mediated 5hmC formation is actively enriched at DNA damage sites [55]. These demethylated cytosine analogues are converted back to cytosine by the base excision repair pathway, thus supporting a role for TET proteins in eliciting the DNA damage response pathway [71, 72]. Conversely, loss of TET suppresses DNA damage response and reduces genomic instability. Our previous work revealed a role for TET proteins in RF stability and indicated that loss of TET2 results in RF protection in BRCA2-deficient cells and confers PARPi resistance [35]. Loss of TET1 suppresses apoptosis and radiation-induced DNA damage response and allows cells to proliferate in the presence of genomic instability [72]. Loss of TET3 leads to reduction in ATR-dependent DNA damage response [73]. Tumor hypoxia causes genomic hypermethylation by reducing the activity of TET proteins, which promotes breast cancer progression and confers selective advantage and, possibly, chemoresistance in tumors [74, 75].

VitC is a co-factor for the α-KDGG class of enzymes and is known to enhance TET activity. VitC has been shown to boost 5hmC levels in a TET-dependent manner in cultured cells and mice [76]. Our mass spectrometry analysis revealed a VitC-dependent increase in 5hmC levels in PARPi-sensitive and PARPi-resistant cells. Certain PARP inhibitors, such as olaparib and talazoparib, trap PARP on the chromatin and selectively kill BRCA1/2-deficient cells. The direct incorporation of 5hmC analogue in the DNA led to an increase in PARP1 trapping when cells were treated with olaparib. Interestingly, Brabson et al. have reported a VitC-dependent increase in TET activity to enhance PARPi cytotoxicity in AML by enhancing PARP trapping [50]. However, they showed that the trapping was due to 5fC [50]. We observed only a mild increase in PARP trapping due to 5fC, and its incorporation had no effect on olaparib sensitivity (Figure 3D-E). The cause of the conflicting impact of 5hmC and 5fC and their role in PARP trapping remains unclear.

In this study, we have shown a previously unknown role of 5hmC and VitC in restoring sensitivity to PARPi in chemoresistant BRCA1/2-deficient cells. Effect of 5hmC on PARPi sensitivity was synergistic, however that of VitC was additive. Also, VitC affected viability of BRCA1/2 deficient cells in absence of PARPi, which suggests that the effect of VitC on BRCA1/2 deficient cells is partially independent of PARP trapping. A high dose of VitC is known to generate oxidative stress and DNA damage [77–79]. Although we did not detect a significant change in ROS levels, we observed a mild increase in DNA damage in VitC-treated cells (Figure 2A-B, 4G-H). Thus, the mechanism behind VitC dependent sensitivity of BRCA1/2 cells remain unclear.

PARP inhibition generates replication gaps that are detrimental for HR-deficient cells [63, 65]. We observed that replication gaps were significantly higher in cells treated with both olaparib and 5hmC or VitC. We propose that excessive PARP1 trapping in response to combined treatment with olaparib and 5hmC/VitC generates DNA lesions that perturb DNA replication, resulting in more replication gaps (Figure 6). Thus, the combined treatment of 5hmC/VitC with olaparib elevates genomic toxicity in the form of replication gaps restoring PARPi sensitivity in chemoresistant cells. It has been shown that replication gaps are a key determinant for PARP sensitivity [65]. Thus, given the clinical significance of PARP inhibitors, seeking drug combinations that maximize replication gaps are likely to enhance PARPi efficacy. Our study has identified the combination of PARP inhibitors and 5hmC or VitC to increase replication gaps in chemoresistant BRCA1/2 deficient cells, when resistance is acquired by restoration of HR function by loss of 53BP1, or stalled replication forks are protected by loss of PTIP or CHD4. When resistance is acquired by secondary mutations that restore BRCA1 or BRCA2 function, 5hmC or VitC is not likely to render cells sensitive to PARPi. In contrast, when resistance is due to lack of PARP expression or overexpression of ABC transporters, 5hmC or VitC treatment can result in hypersensitivity because the cells are deficient in BRCA function. Whether, PARPi will have additional effect in the absence of PARP proteins remains to be tested. Future studies will be focused on testing the effect of 5hmC/VitC in overcoming PARPi resistance under these conditions.

In conclusion, our study demonstrates that a combination treatment of olaparib and 5hmC/VitC is effective against BRCA1/2-deficient chemoresistant cells. This restoration of PARP inhibitor sensitivity is attributed to the ability of 5hmC and VitC to trap PARP1 and increase DNA damage via the formation of replication gaps. Thus, this treatment can be beneficial for patients with cancers resistant to current FDA-approved PARP inhibitor therapies.

### Experimental Procedures

#### Plasmids, Antibodies and Reagents

Western blotting, DNA fiber assays used the following antibodies: PARP1 (9542S; Cell Signaling Technology), Histone H3 (9715S; Cell Signaling Technology), CldU (ab6326; Abcam, 1:500 dilution for DNA fiber assays), IdU (347580; BD Biosciences, 1:500 dilution for DNA fiber assays). The following chemicals were purchased: Olaparib (AZD2281, S1060; Selleck Chemicals LLC), Vitamin C (VitC/L-ascorbate A7631; Sigma-Aldrich), 2′-deoxycytidine (Cytidine, D3897; Sigma-Aldrich), 5-methyl-2’-deoxy cytidine (5mC, M295900; Toronto Research Chemicals), 5-(hydroxymethyl)-2’-deoxycytidine (5hmC, H946630; Toronto Research Chemicals), 5-formyl-2’-deoxycytidine (5fC, PY7589; Berry and Associates), 2’-deoxycytidine-5-carboxylic acid, sodium salt (5caC, PY7593; Berry and Associates).

#### Transient knockdown of proteins

Transient knockdown of proteins was carried out by siRNA transfection using Lipofectamine RNAiMAX transfection reagent (Life Technologies). All siRNAs were purchased as SMARTpool siGENOME category from Dharmacon. Mouse TET1 (L-062861-01-0005), Mouse TET2 (L-058965-00-0005), mouse TET3 (L-054156-01-0005), Human BRCA1 (M-003461-02-0005), Human BRCA2 (M-003462-01-0005), Human 53BP1 (M-003548-01-0005), Human CHD4 (M-009774-01-0005), Human PTIP (M-012795-01-0005). siRNA transfections were performed using Lipofectamine RNAiMAX (Life Technologies) according to manufacturer’s protocol.

#### Cell culture and drug treatments

UWB1.289-WT/BRCA1KO cells cultured in 50% RPMI media, 50% MEGM BulletKit (Lonza CC-3150) supplemented with 3% FBS, 100 U/mL penicillin, and 100 μg/mL streptomycin at 37°C, 5% CO_2_. DLD1-WT/BRCA2KO cells were cultured at 37°C, 5% CO2 in DMEM/F-12 (Life Technologies) supplemented with 10% FBS, antibiotics. DLD1-BRCA2;TET2 KO cells were generated using published CRISPR-Cas9 protocol [80]. BRCA2 deficient Mouse mammary tumor cell line (KB2P1.21) was cultured at 37 °C, 5% CO2, 3% O2 in DMEM/F-12 (Life Technologies) supplemented with 10% FBS, antibiotics. Mouse embryonic fibroblasts MEF-WT/ *Brca1*Δ*11*/ *Brca1*Δ*11;53Bp1KO* were kind gift from Dr. Andre Nussenzweig (National Cancer Institute, Bethesda). MEFs were cultured at 37°C, 5% CO2 in DMEM/F-12 (Life Technologies) supplemented with 10% FBS, antibiotics. Combined treatments for Olaparib and VitC/Cyt/5mC/5hmC/5fC/5caC in clonogenic and XTT cell survival assays were given simultaneously. Treatments for VitC/Cyt/5mC/5hmC/5fC/5caC (48 hrs) and olaparib (10 mM, 2 hrs) in PARP trapping experiments were given sequentially.

#### Cell survival assays

Sensitivity of cells to indicated drug was measured by plating 250-1000 cells per well in 96-well plates in 8 technical replicates for XTT assay and 1000-2500 cells per well for clonogenic assay. Each experiment was repeated 2-3 times as independent biological replicate for each group. The next day, cells were treated with increasing doses of drugs as indicated in corresponding figures and maintained in complete media for 5 to 7 days for XTT assay and 10-12 days for clonogenic assay. Fresh media containing indicated drugs was replenished every 48 hrs. XTT based cell viability was measure by assay as described previously [81]. Colony forming units were stained using 0.5% (w/v) crystal violet in methanol in clonogenic assay.

#### DNA fiber assay

Cells were treated with indicated combinations of olaparib (100 nM), VitC (1 mM) and 5hmC (1 mM) for 48 hrs. After PBS wash cells were labeled by sequential incorporation of two different thymidine analogs, CldU and IdU, into nascent DNA strands for 30 mins. After thymidine analogs were incorporated in vivo, the cells were processed for generation of glass slides with DNA spreads. Briefly, cells were trypsinized, washed with PBS, spotted, and lysed on positively charged microscope slides for 8 min at room temperature. For experiments with the ssDNA-specific endonuclease S1, after the CldU pulse, cells were treated with CSK100 buffer for 10 min at room temperature, then incubated with S1 nuclease buffer with or without 20 U/mL S1 nuclease (Invitrogen, 18001-016) for 30 min at 37°C. The cells were then scraped in PBS + 0.1% BSA and centrifuged at 5,000 rpm for 5 min at 4°C. DNA fibers spreads were generated by tilting the slides at 45°C. After air-drying, fibers were fixed by 3:1 methanol/acetic acid at 4°C overnight. After fixation fibers were rehydrated in PBS, denatured with 2.5 M HCl for 60 min, washed 3 times with PBS for 5mins and blocked with blocking buffer (PBS + 0.1% Triton + 3%BSA) for 40 mins. Next, slides were incubated for 2 hr with primary antibodies for (CldU, Abcam 6326; IdU, Becton Dickinson 347580) diluted in blocking buffer (1:500), washed 3 times in PBS-T (Tween 20), and then incubated with secondary antibodies (CldU, goat anti-rat, Alexa Fluor 594; IdU, goat anti-mouse, Alexa Fluor 488) in blocking buffer (1:1000) for 1 hr. After washing and air-drying, slides were mounted with Prolong (Invitrogen, P36930). The images were captured in Zeiss Axio Imager Z1 microscope, and the fiber length was measured by ImageJ software (NIH). All DNA fiber analysis were performed blindly.

#### Quantitative PCR

qPCR to determine TET1, TET2, TET3, CHD4, PTIP, 53BP1, BRCA1 and BRCA2 mRNA levels in mentioned cell line was performed by using iTaq Universal SYBR Green Supermix (Bio-Rad). qPCR reaction was run on Mx3000P qPCR system (Agilent Technologies).

ROS Assay. ROS levels were measured using a Cellular ROS Assay Kit (Abcam; #ab186027), performed according to manufacturer’s instructions. Briefly, cells were seeded in 96-well plates in 4 technical replicates. After olaparib (100 nM) and VitC (1 mM) treatments for 48 hrs, cells were incubated with ROS Red Stain working solution provided in the kit for 1 h prior to treatment. Treatment solutions made at 10× concentration was then added to the wells to achieve the appropriate treatment concentrations. After a 2 h incubation, fluorescence was measured at 520 nm/605 nm using a microplate reader (SpectraMax iD5, Molecular Devices). ROS measurements were normalized to that of the DMSO condition.

#### Chromatin fractionation and PARP trapping

Cells were treated with gradient or indicated concentrations of VitC or Cyt/5mC/5hmC/5fC/5caC in 10 cm plates for 48 hrs, followed by 2 hr 10 μM olaparib treatment. 1*10^6^ cells were fractionated using a Subcellular protein fractionation kit for cultured cells (ThermoFisher, 78840) according to the manufacturer’s recommendations. PARP1 (9542S; Cell Signaling Technology), Histone H3 (9715S; Cell Signaling Technology) antibodies were used to detect soluble and chromatin bound PARP1 and H3 were detected using Western blotting.

#### Mass spectrometry-based quantitation of cytosine, 5mC and 5hmC

Genomic DNA was isolated by Phenol:chloroform method. 5 mg of genomic DNA was digested to nucleoside level by using Nucleoside digestion mix (M0649S; New England Biolabs). Experiments were repeated 2 times. Each biological replicate contained 3 technical replicates of each sample. 3-5 reactions of digested genomic DNA of each condition was pooled and processed for Mass spectrometry-based quantitation of cytosine, 5mC and 5hmC was performed as described [82, 83]. Briefly, Cytosine, 5mC, 5hmC were quantitated using a Thermo Vanquish Ultra-High-Performance Liquid Chromatography (UHPLC) coupled to a Thermo TSQ-Altis tandem mass spectrometer through an electrospray ion source operating in positive ion mode at 3.5kV. Stock standard solutions were prepared in deionized water at a concentration of 1 mmol/L each. Calibration standard mixtures were prepared at concentrations between 1.0-250umol/L for dC, 0.04-10 umol/L for mdC, and 0.002-0.5 umol/L for hmdC. Linear calibration plots were prepared using concentration versus peak area integration (zero intercept) with a R^2^ greater than 0.999. By comparing the internal standard normalized peak areas in the digest sample to the corresponding retention times from the calibration standards, the micromolar concentrations of the nucleosides were determined against the standard curve. The molar ratio as a percent was then calculated using the formula: <u>Mol% hmdC</u> = ([hmdC] / ([dC] + [mdC] + [hmdC]) × 100

#### ChIP-ELISA

For ChIP-ELISA, BRCA2 deficient DLD1 cells were treated with LAA (250μM) or Olaparib (1μM) or combination of both for 48 hrs. 2X10^6^ cells were crosslinked with formaldehyde and chromatin fragments (200-700bp) were obtained by sonication (Diagenode). PARP1 on the DNA fragments was immunoprecipitated using 3μg of anti-PARP antibody (CST) and the antibody bound fractions were recovered with Protein-G and A dynabeads overnight. Next day, the samples were washed, eluted and the DNA was purified using EZ-Magna ChIP™ A/G Chromatin Immunoprecipitation Kit as per manufacturers protocol (Millipore). For quantifying, methylcytosine and oxidized methylcytosine in ChIP DNA samples, 5mC, 5-hmC, and 5-fC ELISA kit (Epigentek) were used. 30ng DNA were used to coat the ELISA plate and the experiment was carried out as per manufacturers protocol. Samples were analyzed in technical duplicates.

#### Statistics

Statistics analysis of cell survival assays was performed using paired T test. Statistical analysis of DNA fiber assays and quantitation of foci was performed by using two-tailed *t*-test, Mann–Whitney test. All error bars represent standard deviation. *P*<0.05 was considered statistically significant: ^ns^ *P*≥0.05, * *P* ≤0.05, ** *P* ≤ 0.01, *** *P*≤0.001, and **** *P*≤0.0001. No statistical methods or criteria were used to estimate sample size or to include or exclude samples. The investigators were not blinded to the group allocation during the experiments.

## Supporting Information

This article contains supporting information.

## Acknowledgements

We thank all the members of our laboratory for their helpful suggestions. We thank Sounak Sahu for critical review of the manuscript. We thank Drs. Andre Nussenzweig and Elsa Callén (NCI, Bethesda) for providing BRCA1/53BP1 MEFs and Dr. Yves Pommier and Dr. Yilun Sun (NCI, Bethesda) for help with PARP trapping experiments.

## Author contribution

S.S.K. performed most of the experiments and analyzed the data. A.P.M. performed IF experiment to examine γH2AX foci formation and helped with PARP trapping studies. S.K.Sengodan performed ChIP-ELISA. D.D helped with DNA fiber assay. S.D.F. performed mass spectrometric analysis, S.K.Sharan conceived and supervised the study. S.S.K. and S.K.Sharan. wrote the manuscript. All the authors reviewed and commented on the manuscript.

## Funding and additional information

This research was sponsored by the Intramural Research Program, Center for Cancer Research, National Cancer Institute, US National Institutes of Health. The content is solely the responsibility of the authors and does not necessarily represent the official views of the National Institutes of Health.

