## Supplementary Figures 1-5 for "5hmC enhances PARP trapping and restores PARP inhibitor sensitivity in chemoresistant BRCA1/2-deficient cells"

### Supplementary Figure 1 – VitC enhances PARP inhibitor sensitivity in BRCA1/2 deficient cells

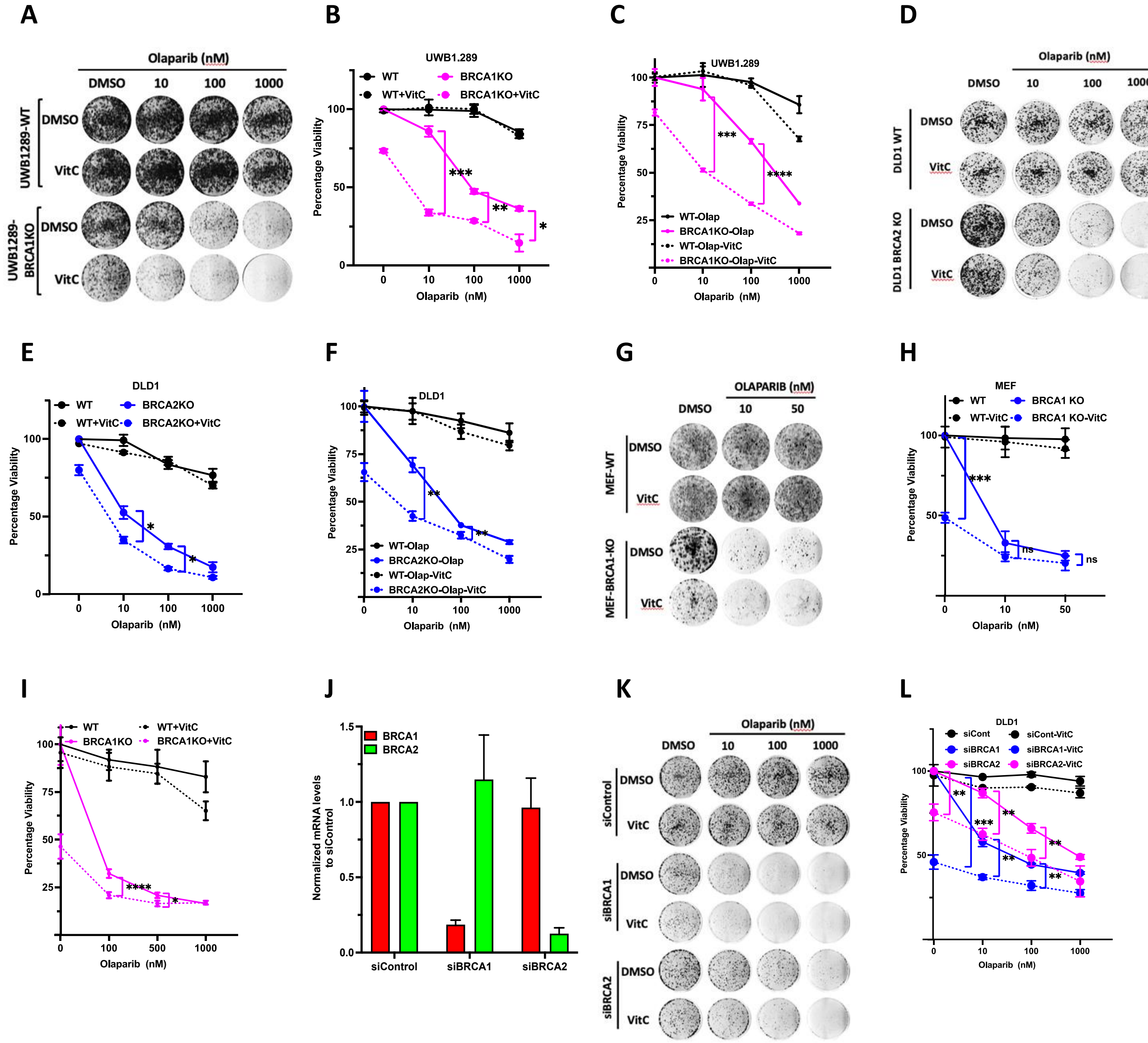

### Supplementary Figure 2 – VitC/TET dependent 5hmC increase enhances PARP inhibitor sensitivity in BRCA1/2 deficient cells

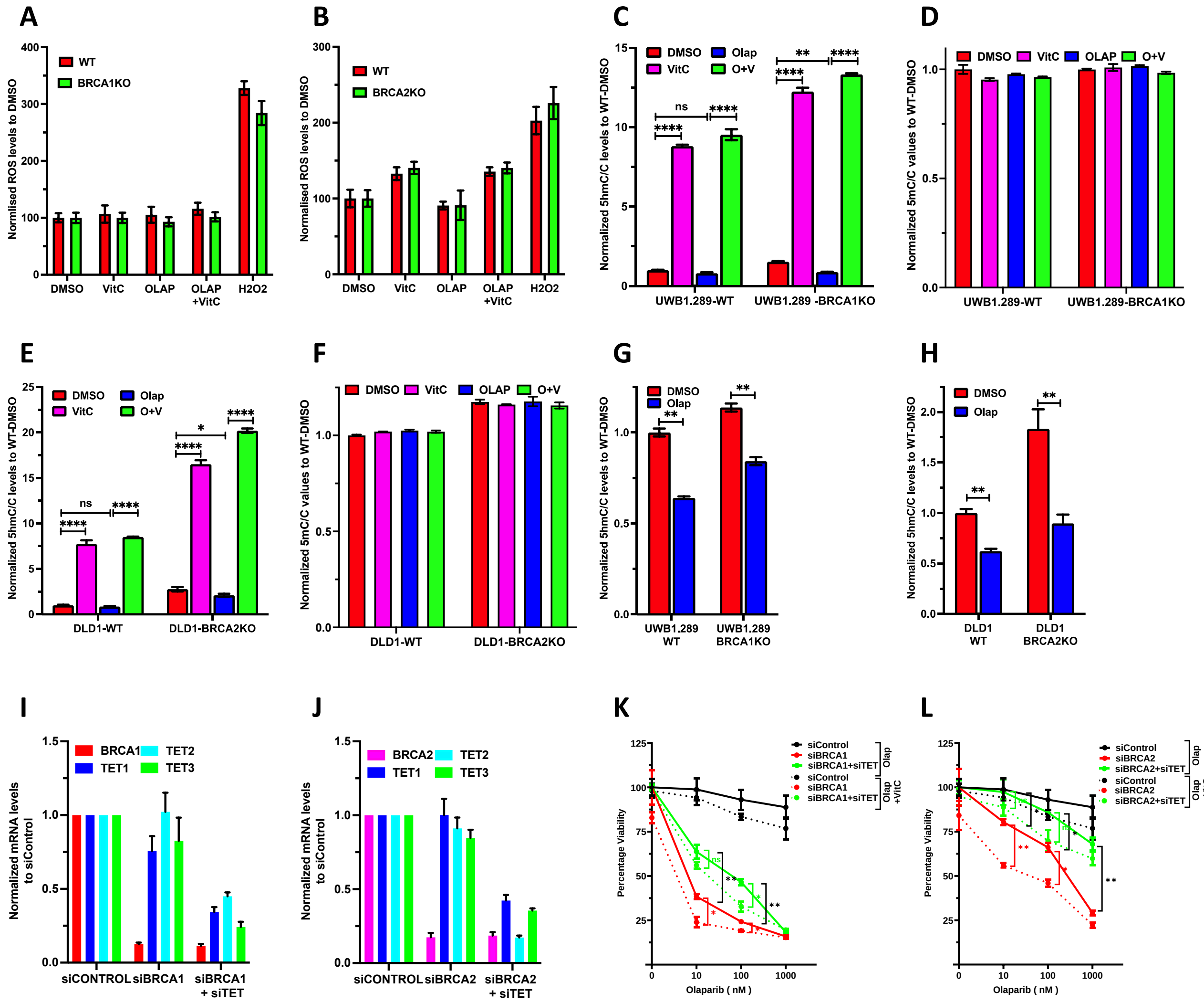

### Supplementary Figure 3 – VitC and 5hmC restores PARP inhibitor sensitivity in chemoresistant BRCA1/2 deficient cells

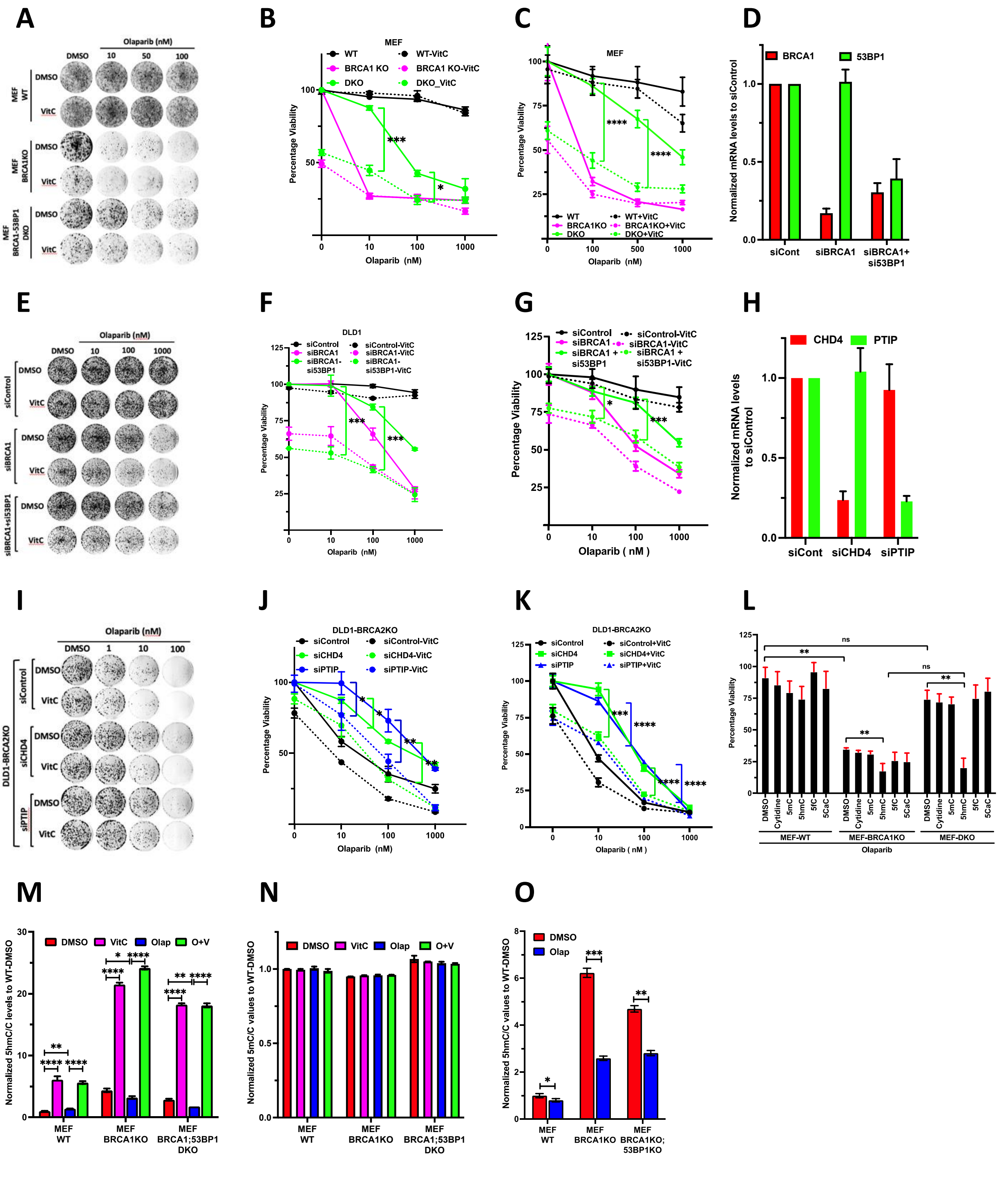

#### Supplementary Figure 4 – VitC traps PARP1 in presence of olaparib on chromatin and increase DNA damage in BRCA1/2 deficient cells

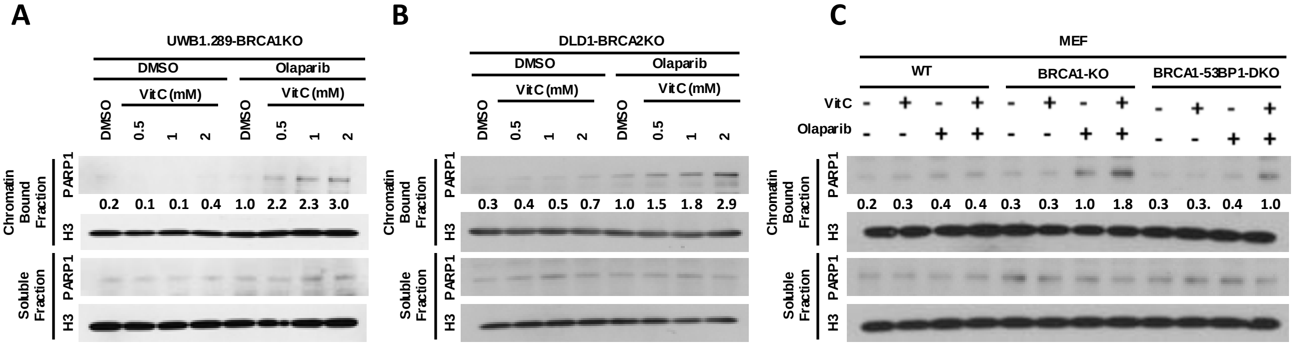

### Supplementary Figure 5 – VitC increases replication gaps and DNA damage in olaparib treated BRCA1/2 deficient cells

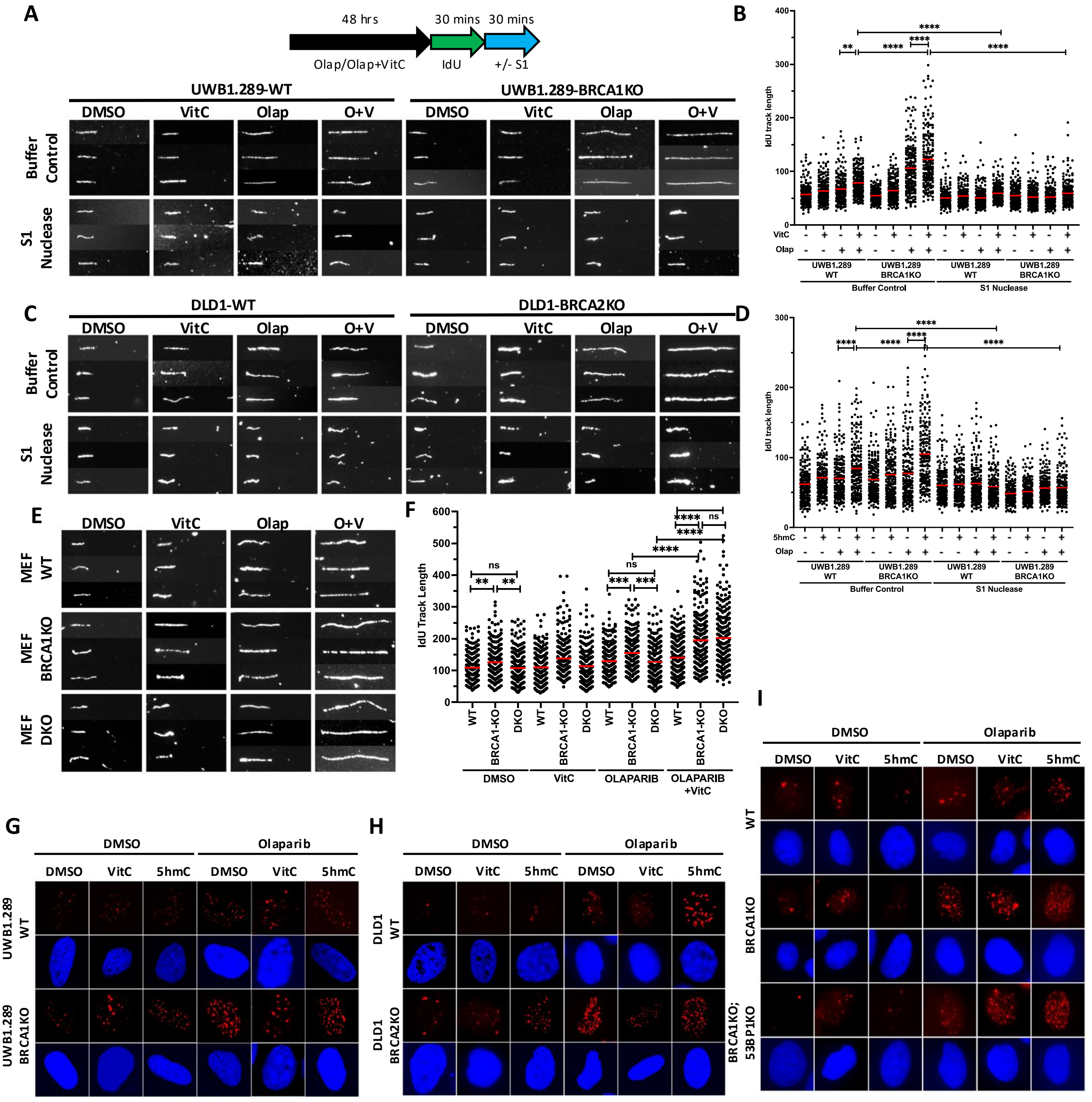
